# Altered population activity and local tuning heterogeneity in auditory cortex of *Cacna2d3*-deficient mice

**DOI:** 10.1101/2022.08.30.505788

**Authors:** Simon L. Wadle, Tatjana T.X. Schmitt, Jutta Engel, Simone Kurt, Jan J. Hirtz

## Abstract

The α_2_δ3 auxiliary subunit of voltage-activated calcium channels is required for normal synaptic transmission and precise temporal processing of sounds in the auditory brainstem. In mice its loss additionally leads to an inability to distinguish amplitude-modulated tones. Furthermore, loss of function of α_2_δ3 has been associated with autism spectrum disorder in humans. To investigate possible alterations of network activity in the higher-order auditory system in α_2_δ3 knockout mice, we analyzed neuronal activity patterns and topography of frequency tuning within networks of the auditory cortex (AC) using two-photon Ca^2+^ imaging. Compared to wild-type mice we found distinct subfield-specific alterations in the primary auditory cortex, expressed in overall lower correlations between the network activity patterns in response to different sounds as well as lower reliability of these patterns upon repetitions of the same sound. Higher AC subfields did not display these alterations but showed a higher amount of well-tuned neurons along with lower local heterogeneity of the neurons’ frequency tuning. Our results provide new insight into AC network activity alterations in an autism spectrum disorder-associated mouse model.

## Introduction

Sound processing within networks of the auditory cortex (AC) is highly complex, featuring not only the perception of sounds (Ohl et al. 1999; Wetzel et al. 2008; Homma et al. 2017; O’Sullivan et al. 2019), but also their contextualization, for example in the process of auditory-related learning (Ono et al. 2006; Bajo et al. 2010; Schulze et al. 2014; Caras and Sanes 2017; Schneider et al. 2018; Xin et al. 2019) or in aspects of communication (Kim and Bao 2013; Tasaka et al. 2018; Moore and Woolley 2019; Zhu et al. 2019; Montes-Lourido et al. 2021). It has been reported that decision making can be decoded from AC population activity (Bathellier et al. 2012; Xin et al. 2019) and that acute AC inactivation using pharmacology or optogenetics disturbs the discrimination of even basic sound features (Talwar et al. 2001; O’Sullivan et al. 2019). These observations provide a strong incentive to study AC network activity in models of central auditory processing defects. It has recently been reported that *Cacna2d3*-deficient (α_2_δ3 KO) mice exhibit an interesting behavioral abnormality, as they are unable to distinguish two amplitude-modulated (AM) tones (20 versus 40 Hz modulation frequency) from each other in an auditory discrimination task, while distinguishing pure tones (PTs) was undisrupted (Pirone et al. 2014). Furthermore, α_2_δ3 KO mice display distorted auditory brainstem responses, despite nearly normal hearing thresholds, as well as malformed and functionally impaired synapses in the auditory brainstem (Pirone et al. 2014). Recordings from neurons in the inferior colliculus (IC) confirmed impaired temporal processing, although strong deficits were only identified above 70 Hz modulation frequency (Bracic et al. 2022).

The neuronal extracellular membrane-anchored α_2_δ proteins 1-3 assist in trafficking and surface expression of the voltage gated Ca^2+^ channel (Ca_V_) complex but some of them fulfil additional functions including channel trafficking along axons, synapse development, and trans-synaptic alignment (Dolphin 2012; Ablinger et al. 2020; Schöpf et al. 2021). The *Cacna2d3* gene encodes for the auxiliary α_2_δ3 voltage gated Ca^2+^ channel subunit (Neely et al. 2010; Dolphin 2012). Importantly, loss of function mutations of this gene have recently been identified as risk mutations for autism spectrum disorder (ASD) in humans (Iossifov et al. 2012; Girirajan et al. 2013; De Rubeis et al. 2014). As auditory deficits have been reported in people with ASD (Robertson and Baron-Cohen 2017; Vlaskamp et al. 2017; Keehn et al. 2019; Schafer et al. 2020), studying activity patterns within the central auditory system of α_2_δ3 KO mice might provide more insight into the underlying sound processing defects.

Prior to 2010, studies of AC network activity in rodents employed almost exclusively electrical recordings. While these provide precise temporal resolution, the introduction of two-photon imaging enabled an unprecedented spatial precision while maintaining single-cell resolution. The works of Bandyopadhyay et al. (2010) and Rothschild et al. (2010), which were corroborated and expanded by further studies using the same technique (Romero et al. 2019; Tischbirek et al. 2019; Gaucher et al. 2020), revealed a high local heterogeneity of frequency tuning on single-cell level, while clear tonotopy is only observed at a global scale. Two-photon imaging has also enabled the identification of large-scale network activity patterns from which the animal’s decision making can be decoded (Bathellier et al. 2012; Xin et al. 2019).

In the present study, we use two-photon imaging to study activity patterns of AC layer 2/3 neurons of α_2_δ3 KO mice and littermate controls. We demonstrate that alterations are present both on the topographical level as well as expressed in lower correlations of network activity in response to specific sounds, and lower reliabilities of these sound patterns.

## Results

To investigate sound-evoked activity patterns of AC neurons in awake α_2_δ3 KO mice and littermate wildtype (WT) controls, we injected AAV1-hSyn-GCaMP7f into the AC and performed activity imaging starting 13 days later. To identify AC subfields, widefield (WF) imaging was performed. In both KO animals and controls, hubs preferring either high or low sound frequencies could be observed (Figure 1A, B), allowing for the parcellation into the primary auditory field (A1), the anterior auditory field (AAF), and the secondary auditory field (A2). In some animals, further parcellation was possible (Figure 1B), resulting in identification of A1, AAF, the suprarhinal auditory field (SRAF), the ventro-posterior auditory field (VAPF), and dorsal posterior field (DP), following the nomenclature in Romero et al. (2019). To streamline further analysis, SRAF was termed A2, and DP and VAPF were included into A1. In addition, adjacent tone responsive regions were merged with the nearby subfield (compare low-frequency hub lateral to AAF in 1B middle vs bottom). While these results demonstrate that α_2_δ3 KO mice display rough-scale tonotopic organization similar to those observed in wild type mice, we next aimed to characterize sound-evoked properties at single-cell level in layer 2/3, performing two-photon imaging. Field-of-views (FOVs) were aligned with WF maps to assign subfield identity to each neuron imaged (Figure 1C).

**Figure 1:**
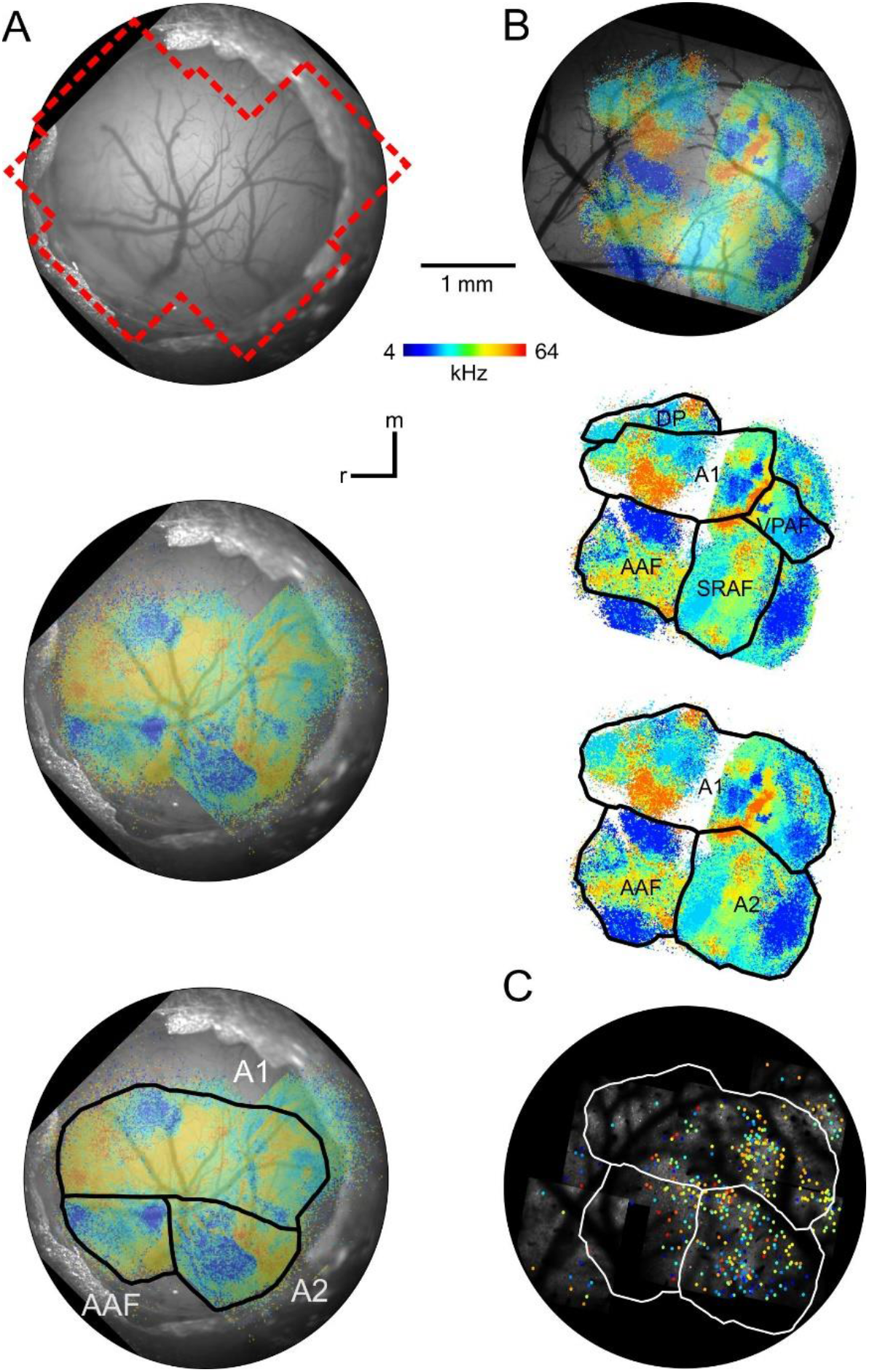
Widefield maps and subfield alignment. (A) Top: Image of a cranial window from a α_2_δ3 KO animal with merged borders of 10x epifluorescence FOVs (red). Middle: Same cranial window as top but with pixel-wise BF color coding based on the fluorescence signal during tone stimulation. Bottom: Same as middle but the three assigned subfield borders are depicted. (B) Top: Superimposed BF false color map of a WT cranial window. Middle: Same BF map as left with borders of five subfields (see main text). Bottom: Same map as middle but with three borders drawn by merging and adding tone responsive regions to the nearest subfield. (C) Alignment of 2P imaging experiments with the WF map presented in B. Colored dots depict BF of single-peak neurons.

First, 17 PTs of 4-64 kHz were played at different sound pressure levels (30-70 dB SPL, 10 dB steps, Supplementary Figure 1). After region of interest (ROI) extraction and deconvolution of neuronal activity from Ca^2+^ traces, a frequency response area (FRA) was created for each neuron recorded. Only neurons with a significant (one-way ANOVA; p <0.01) response in at least one frequency-intensity combination were used for following analyses and were termed “PT-responsive” (WT: 3746, KO: 4606). Typically, AC neurons are subdivided into three PT response categories, single- and double-peaked and complex (here termed “irregular-tuned”) (Gaucher et al. 2020). Using gaussian fit functions we categorized each neuron, finding that the majority in both genotypes was irregular-tuned followed by single-peaked neurons (WT: 80 % and 16 %, KO: 78 % and 19 %, respectively). Further analysis concentrated mostly on single-peaked neurons, which displayed relatively sharp tuning in both WT (653 neurons) and KO (1074 neurons, Figure 2A and Supplementary Figure 2). Tuning bandwidth was assessed for all WT and α_2_δ3 KO neurons which showed single-peak tuning. It did not differ between genotypes for A1 and A2 (A1: WT: 0.71 oct ± 0.03, KO: 0.66 oct ± 0.03, *p* = 0.32; A2: WT: 0.80 oct ± 0.04, KO: 0.80 oct ± 0.02, *p* = 0.91). However, neurons in AAF of α_2_δ3 KOs had a broader bandwidth compared to WT (WT: 0.71 oct ± 0.06, KO: 0.88 oct ± 0.02, *p* = 0.007). To analyze the representation of frequencies across the populations observed, we next reduced the neurons’ tuning information to their best frequencies (BF). Typically, neurons in A1 show a skewed BF distribution towards mid-frequencies (10-28 kHz, Moore and Wehr 2013; Joachimsthaler et al. 2014; Panniello et al. 2018). We also observed that BF distributions of each subfield in WTs had an overrepresentation of frequencies in the mid-to-high frequency range (16 kHz – 32 kHz, Figure 2B). BF distributions in α_2_δ3 KOs displayed a similar pattern in A1 but showed a reduced high and low frequency representation in A2, with only 1.2 % of single-peak neurons having a BF above 32 kHz (Figure 2B). Furthermore, the BF distribution in AAF was slightly shifted towards lower frequencies (4-8 kHz), representing 42 % of neurons in α_2_δ3 KO versus 22 % in WTs. Notably, this was accompanied by an overrepresentation of single-peaked neurons in AAF and A2, but not in A1 (Figure 2C).

**Figure 2:**
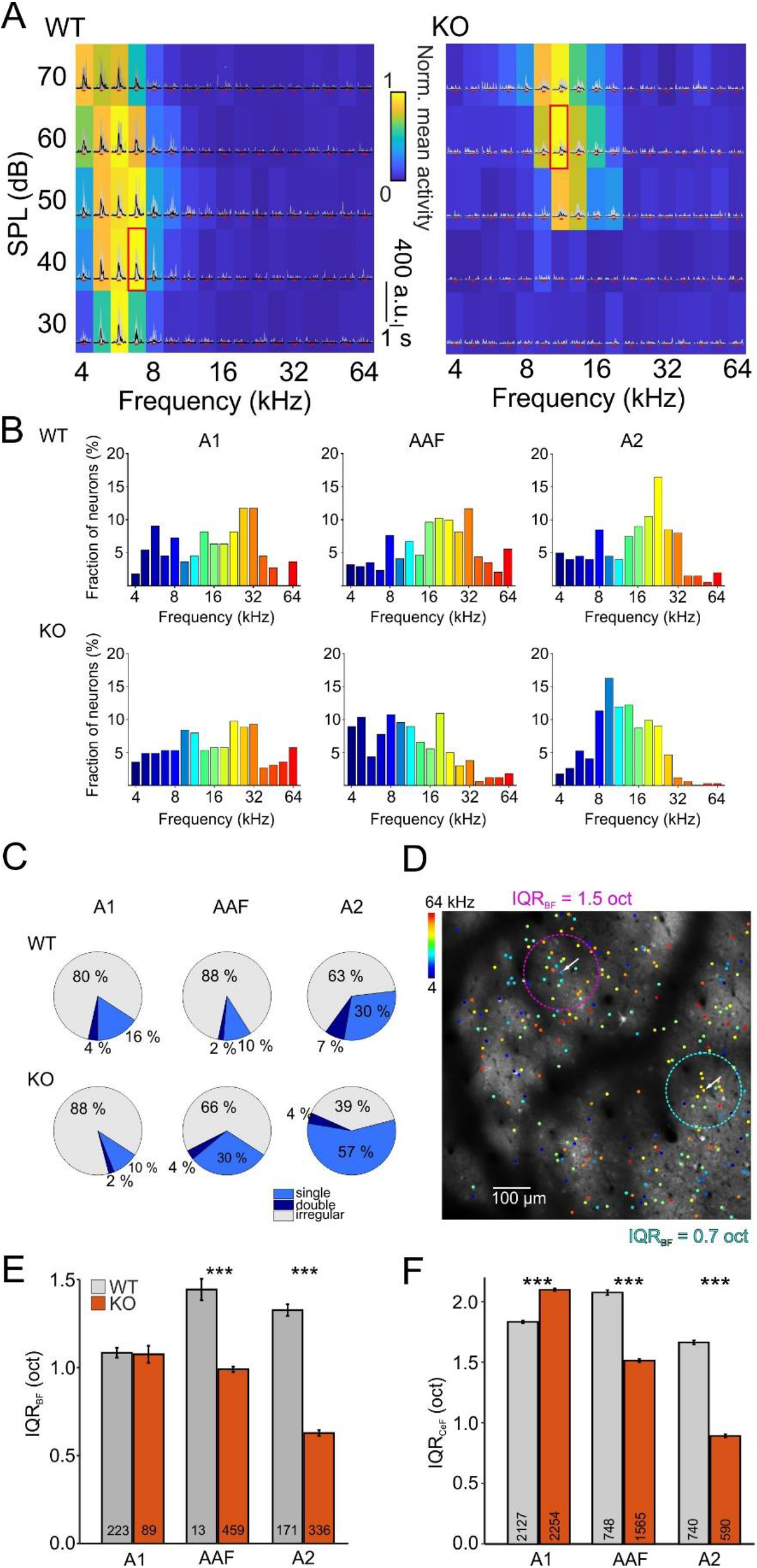
Tuning distribution and local heterogeneity. (A) Sorted and cut activity traces (deconvolution of fluorescence traces) upon PT stimulation of one exemplary neuron from each, a WT and a α2δ3 KO animal, displaying a single-peak tuning. 10 repetitions in grey, mean in black and timing of tone presentation is displayed by red lines below traces. The background depicts color-coded normalized activity (resulting in the FRA). Activity during 400 ms following tone onset was averaged for each frequency-intensity combination and normalized to the maximal response. BF is depicted by a red rectangle. (B) BF distributions of single-peak neurons within three AC subfields in WT and KO. (C) Fraction of single-peaked, double-peaked and irregular-tuned neurons. (D) Two-photon image with superimposed color-coded dots depicting BFs of single-peak neurons. Local IQR calculation is shown for two α2δ3 KO example neurons (white arrows). Cyan and purple dotted circles show the area considered for IQR analysis. (E) Local heterogeneity analysis of BF of all single-peak neurons. (F) Local heterogeneity analysis of CeF of all PT-responsive neurons.

Whereas linear tonotopy on the mesoscale is well established (Guo et al. 2012) quite a high local heterogeneity of tuning has been observed at the single cell level (Bandyopadhyay et al. 2010; Panniello et al. 2018; Romero et al. 2019; Tischbirek et al. 2019; but see Issa et al. 2014). To assess whether the tonotopic order within AC subfields differs between α_2_δ3 KOs and WT controls, we calculated the interquartile range (IQR) of BFs for each neuron within a 100 µm radius, an approach used before for this purpose (Romero et al. 2019; Bowen et al. 2020; Liu and Kanold 2021). Figure 2D shows two α_2_δ3 KO example neurons with an IQR_BF_ of 1.5 oct and 0.7 oct, representing a less and highly organized region, respectively. Interestingly, population data showed a reduced local heterogeneity for α_2_δ3 KOs again in AAF and A2, but not in A1. (A1: WT: 1.08 oct ± 0.03, KO: 1.08 oct ± 0.05, *p* = 0.18; AAF: WT: 1.44 oct ± 0.06, KO: 0.99 oct ± 0.02, *p* = 8.5 × 10^−4^; A2: WT: 1.33 oct ± 0.03, KO: 0.63 oct ± 0.02, *p* = 6.6 × 10^−43^; Figure 2E), suggesting a more precise tonotopic organization in α_2_δ3 KO mice. Furthermore, we aimed to include all PT-responsive neurons in this analysis to not limit our observations to well-tuned neurons, and thus determined the center frequency (CeF), the frequency at which a single Gaussian fit calculated the strongest response of any given neuron. This parameter provides a measure for each neuron’s overall preferred frequency, as, unlike BF, it takes the complete FRA into account. However, it has the caveat of a CeF in the middle of the analyzed spectrum being the result of either specific preference of that frequency, or an overall unspecific response to PTs. Differences in local heterogeneity were similar for CeF compared to BF, with the exception of IQR_CeF_ being slightly higher in A1 of KO mice compared to WT controls (A1: WT: 1.83 oct ± 0.01, KO: 2.1 oct ± 0.01, *p* = 3.8 × 10^−69^; AAF: WT: 2.08 oct ± 0.02, KO: 1.51 oct ± 0.02, *p* = 4.28 × 10^−88^; A2: WT: 1.66 oct ± 0.02, KO: 0.89 oct ± 0.01, *p* = 1.1 × 10^−138^; Figure 2F).

α_2_δ3 is widely expressed in the brain, including cortical areas (Landmann et al. 2018). Thus, a potential concern for using Ca^2+^ imaging as activity readout when using α_2_δ3 KO mice is the involvement of the subunit in Ca_V_ trafficking, expression control and modulation of Ca_V_ biophysical properties (Dolphin 2012; Ablinger et al. 2020; Schöpf et al. 2021). To assess whether this might impair action potential (AP) detection, we performed patch-clamp recordings in acute slices using pipettes containing a calcium indicator. The amplitude of AP-evoked calcium traces was similar between KO mice and WT controls, both for single APs (WT: 3.6 % ± 0.4, KO: 3.0 % ± 0.7, *p* = 0.38, Supplementary Figure 3) as well as peaks of responses evoked at 2 Hz (WT: 8.8 % ± 1.2, KO: 6.6 % ± 1.4, *p* = 0.33), 5 Hz (WT: 11.5 % ± 1.5, KO: 8.8 % ± 1.9, *p* = 0.33) and 10 Hz (WT: 12.4 % ± 1.8, KO: 10.1 % ± 2.2, *p* = 0.53). We thus concluded that although our analysis cannot exclude small differences in Ca_V_-mediated currents between genotypes, these are very unlikely to cause diminished identification of activity in two-photon imaging experiments. Our results are also in line with findings in cultured dissociated spiral ganglion neurons derived from α_2_δ3 KO mice, in which the overall reduction in Ca^2+^ currents is minimal if existent at all, though currents of specific Ca_V_ types are strongly affected (Stephani et al. 2019).

As noted above, α_2_δ3 KO mice failed to distinguish AM tones with different modulation frequencies in a behavioral task. To investigate whether this might be due to impaired AC processing, we presented AM tone sets with either 20 or 40 Hz modulation frequency as well as PTs of the same carrier frequencies (17 tones at 4-64 kHz per set). Hierarchical clustering of correlations between sound-evoked network activity patterns revealed overall worse correlations within A1 in KO mice as well as worse mean correlation per cluster, a smaller number of clustered sounds, and fewer sounds per cluster, while the number of clusters was similar (Table 1, Figure 3 A, B, and Supplementary Table 1). Furthermore, the reliability of the network responses to sounds upon repetitions (correlation depicted in diagonal in correlation matrix) was lower. While this might imply that categorization of sounds within A1 is worse in KO mice, which would be in line with the described learning deficit, we found this not to be the case. When determining the sound category (either PTs, 20 Hz modulated AM tones, or 40 Hz modulated AM tones) dominating each given cluster, its fraction for each given cluster was similar for KO mice and WT controls (Table 1 and Figure 3B right). The fraction of clusters dominated by either of the sound categories was also quite similar (WT: 25 % PTs, 35 % 20 Hz modulated AM tones, 40 % 40 Hz modulated AM tones; KO: 32 % PTs, 32 % 20 Hz modulated AM tones, 36 % 40 Hz modulated AM tones). This implies that rather than failing to identify sound categories, processing impairments in KO mice are manifested in unreliable network activity patterns and generally weak correlations between sounds. This is not limited to AM and pure tones, as repeating the experiments using ten spectrally complex animal vocalizations yielded similar results concerning correlation, reliability, fraction of clustered sounds, and sounds per cluster (Table 2, Figure 3C, D, and Supplementary Table 2). Interestingly, these impairments were limited to A1 and not observed within AAF or A2 (Tables 1, 2, Supplementary Figure 4, and Supplementary Tables 1, 2), although the number of analyzed FOVs within A2 should be increased to allow for a definitive statement.

**Table 1:** Sound clusters in relation to AM/PT stimulation Features of hierarchically clustered activity patterns evoked by AM tones and PTs in WT and KO mice. The data are also presented in Figure 3 and Supplementary Figure 4.

|  | Overall correlation | Correlation within clusters | Number of clusters | Sounds per cluster | Clustered fraction | Overall reliability | Cluster dominance |
| --- | --- | --- | --- | --- | --- | --- | --- |
| A1 WT | 0.227 ± 0.019 | 0.268 ± 0.01 | 3.9 ± 0.7 | 10.1 ± 1.5 | 0.77 ± 0.05 | 0.248 ± 0.018 | 0.72 ± 0.03 |
| A1 KO | 0.173 ± 0.015 | 0.224 ± 0.008 | 5.9 ± 0.7 | 4.5 ± 0.5 | 0.52 ± 0.05 | 0.183 ± 0.016 | 0.77 ± 0.03 |
| *p*-value | 0.04 | 5.25e-04 | 0.053 | 3.98e-04 | 5.47e-04 | 0.016 | 0.23 |
| AAF WT | 0.227 ± 0.016 | 0.278 ± 0.009 | 5.3 ± 0.6 | 6.0 ± 0.7 | 0.62 ± 0.04 | 0.251 ± 0.017 | 0.74 ± 0.02 |
| AAF KO | 0.225 ± 0.02 | 0.26 ± 0.01 | 5.9 ± 0.9 | 5.6 ± 0.8 | 0.65 ± 0.05 | 0.247 ± 0.022 | 0.75 ± 0.03 |
| *p*-value | 0.95 | 0.21 | 0.6 | 0.5 | 0.7 | 0.868 | 0.67 |
| A2 WT | 0.168 ± 0.032 | 0.23 ± 0.015 | 6.5 ± 1.1 | 5.5 ± 0.9 | 0.7 ± 0.06 | 0.207 ± 0.035 | 0.72 ± 0.03 |
| A2 KO | 0.183 ± 0.043 | 0.242 ± 0.016 | 6.8 ± 1.6 | 5.6 ± 1.1 | 0.74 ± 0.07 | 0.235 ± 0.055 | 0.76 ± 0.04 |
| *p*-value | 0.78 | 0.62 | 0.88 | 0.81 | 0.66 | 0.67 | 0.33 |

**Table 2:** Sound clusters in relation to animal vocalizations Features of hierarchically clustered activity patterns evoked by complex sounds in WT and KO mice. The data are also presented in Figure 3 and Supplementary Figure 4.

|  | Overall correlation | Correlation in clusters | Number of clusters | Sounds per cluster | Clustered fraction | Overall reliability |
| --- | --- | --- | --- | --- | --- | --- |
| A1 WT | 0.332 ± 0.023 | 0.358 ± 0.021 | 1.3 ± 0.1 | 6.7 ± 0.6 | 0.87 ± 0.02 | 0.34 ± 0.02 |
| A1 KO | 0.228 ± 0.02 | 0.274 ± 0.013 | 1.6 ± 0.2 | 4.5 ± 0.5 | 0.7 ± 0.07 | 0.23 ± 0.02 |
| *p*-value | 0.0017 | 9.2e-04 | 0.15 | 0.005 | 0.0073 | 0.0015 |
| AAF WT | 0.28 ± 0.021 | 0.351 ± 0.019 | 1.8 ± 0.2 | 4.1 ± 0.4 | 0.74 ± 0.04 | 0.28 ± 0.02 |
| AAF KO | 0.275 ± 0.02 | 0.33 ± 0.014 | 1.6 ± 0.2 | 5.1 ± 0.5 | 0.8 ± 0.05 | 0.28 ± 0.02 |
| *p*-value | 0.88 | 0.4 | 0.16 | 0.22 | 0.11 | 0.89 |
| A2 WT | 0.252 ± 0.053 | 0.363 ± 0.026 | 2.4 ± 0.3 | 3.5 ± 0.5 | 0.83 ± 0.05 | 0.25 ± 0.06 |
| A2 KO | 0.273 ± 0.039 | 0.339 ± 0.029 | 2.1 ± 0.4 | 3.9 ± 0.6 | 0.82 ± 0.05 | 0.28 ± 0.04 |
| *p*-value | 0.76 | 0.54 | 0.59 | 0.57 | 0.97 | 0.73 |

**Figure 3:**
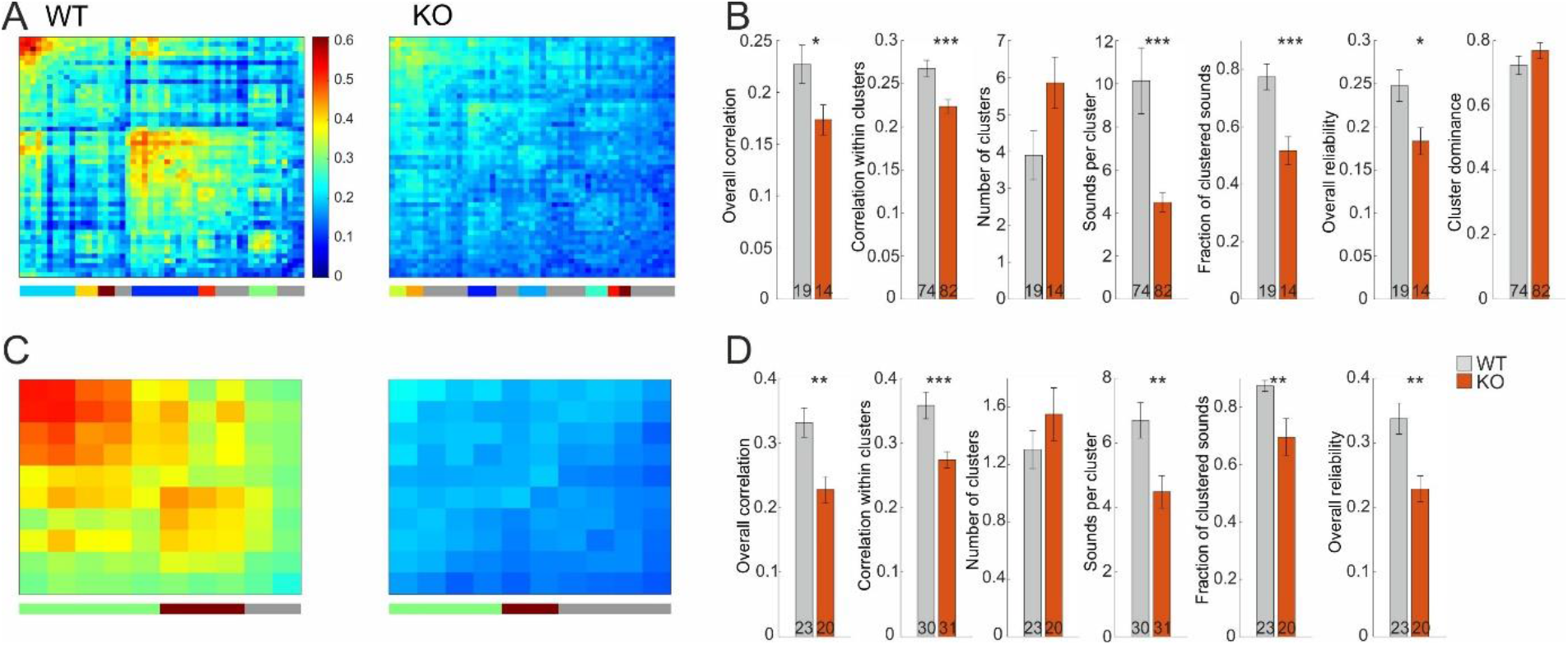
Cluster analysis of sound-evoked activity in A1. (A) Correlation matrix of 51 sound-evoked patterns (17 patterns for each of PTs, AM tones with 20Hz modulation, and AM tones with 40 Hz modulation) after hierarchical clustering. Color code depicts correlation value. Vertical color bar at the bottom depicts clusters, grey stripes correspond to sounds that are not part of clusters. The diagonal depicts mean correlation across repetitions, thus showing reliability of the network. (B) Statistics for sound cluster analysis of PT/AM sounds. Numbers in bars depict n-number, either FOVs or sound clusters. (C) As in A, but for 10 natural animal vocalizations used as sound stimulation. (D) As in B, but for animal vocalizations. See Supplementary Tables 1 and 2 for obtained order of sounds.

Overall, our results show that α_2_δ3 KO mice display alterations in sound processing within AC networks. For A1 this is manifested in generally weaker correlations and reliability of sound-evoked responses, while AAF and A2 show alterations in the topography and tuning distribution when mapping PT-evoked responses.

## Discussion

We have investigated network activity in the AC of α_2_δ3 KO mice and found two types of alterations depending on the cortical subfield. For A1, local heterogeneity of the neurons’ frequency tuning was unaltered to slightly increased, depending on the subset of neurons analyzed, along with an unaltered distribution of BFs and a similar fraction of well-tuned neurons compared to WT controls. In contrast, population activity patterns in response to a wide variety of sounds were strongly affected, displaying weaker correlations between sounds as well as weaker reliability upon repetitions of the same sound. Thus, clustering of sound-specific network patterns resulted in a lower fraction of clustered sounds as well as fewer sounds per clusters, and a slight tendency of more clusters overall. While these network patterns appeared to be unaltered in AAF and A2, local frequency tuning was affected in these subfields, expressed in a lower heterogeneity of BFs around a 100 µm radius of a given neuron. This change might be explained by a less broad distribution of BFs overall, resulting in lower frequency tuning heterogeneity within the complete population. Furthermore, the fraction of well-tuned neurons was higher for KO mice compared to WT controls in AAF and A2.

These alterations are probably a result of impaired fine structure of temporal processing in the auditory brainstem, as Bracic et al. (2022) found for AM tones in neurons of the IC. This in turn might be caused by impaired synaptic transmission in the auditory brainstem, as observed at the endbulb of held in the cochlear nucleus (Pirone et al. 2014), which is in line with α_2_δ subunits being involved in organization of glutamatergic synapses (Schöpf et al. 2021). In addition to impaired temporal processing, a higher rate of spontaneous activity together with lower sound-evoked rates in the IC (Bracic et al. 2022) might be another factor for impaired AC processing, as the signal to noise ratio passed on is decreased. The fact that the AC reacts to this altered input with pronounced, yet not massive activity pattern changes highlights the high compensatory plasticity and adaptability of cortical networks. Obviously, direct network alterations within AC are possible as well, such as an imbalance of excitation and inhibition associated with loss or expression alterations of α_2_δ3 (Landmann et al. 2018; Bikbaev et al. 2020). Expression patterns of α_2_δ subunits are heterogeneous throughout the brain, yet α_2_δ3 mRNA is highly expressed in regions involved in auditory information processing and somatic movement, including the cortex (Cole et al., 2005). However, compensatory effects of other α_2_δ subunits are to be expected, as has been recently elegantly shown in α_2_δ1/α_2_δ2/α_2_δ3 single and double knockout studies (Geisler et al., 2021). This probably offsets overall Ca^2+^ influx, as seen by Stephani et al. (2019), and also suggested by our control experiments. However, the prominent deficits of α_2_δ3 KO mice in the auditory and somatosensory pathway (Pirone et al. 2014, Neely et al. 2010, Landmann et al. 2018, Stephani et al. 2019, Bracic et al. 2022) clearly indicate specific functions of the α_2_δ3 protein. Presumably, Ca_V_-independent functions of a α_2_δ3, e.g. in synaptogenesis, cannot be compensated in this context (Kurshan et al. 2009). Furthermore, α_2_δ3 KO mice also exhibit abnormal cross-modal sensory activation (Neely et al. 2010; Landmann et al. 2018; Landmann et al. 2019), which somewhat complicates drawing final conclusions about the cause of AC activity alterations. Nevertheless, α_2_δ3 KO mice provide an interesting model to study the effects of aberrant central auditory processing, especially because their inner ear functions and hearing thresholds appear to be almost unaffected (Pirone et al. 2014).

The inability of KO mice to distinguish AM tones with 20 or 40 Hz modulation from each other (Pirone et al. 2014) is in line with the observation that for modulations at 30 Hz or higher, phasic neurons in the IC respond slower, yet still is somewhat surprising, as complete failure of phase locking in the IC only occurs at 70 Hz or higher (Bracic et al. 2022). Our findings confirm that AM tones at 20 and 40 Hz modulation can still be distinguished within AC networks, as activity clusters are mostly dominated by one of the two sound categories (or the third tested category, being PTs). However, a lower fraction of sound-evoked activity patterns contributing to clustered activity and decreased reliability upon repetitions in A1 indicate overall less robust and weakly coordinated activity, which might be the underlying cause of the learning defect. Furthermore, our data demonstrate that these network activity alterations are not limited to the rather ‘artificial’ AM tones but observed as well when presenting natural animal vocalizations. The fact that no such alterations were found within AAF or A2 points towards specific roles of AC subfields for auditory learning. It has been shown that task-relevant sounds increase activity levels in A2 stronger than in A1 (Atiani et al. 2014). Furthermore, projections from the higher-order auditory thalamus to the AC, which are involved in associative memory, are more pronounced in A2 than A1 (Pardi et al. 2020). Nevertheless, our results suggest that A1 is highly important when learning to distinguish sounds from each other, which is in line with its pharmacological or optogenetic inhibition leading to impaired sound discrimination (Talwar et al. 2001; O’Sullivan et al. 2019). One might speculate that A1 is more important in deciphering sound structure, and higher auditory fields for decision making, and thus both must be functional for correct behavior in a sound discrimination task. However, these conclusions should be verified by two experiments in the future: i) increasing the number of FOVs analyzed in A2 in α_2_δ3 KO mice and WT controls, and ii) performing auditory-related learning experiments while simultaneously observing AC network activity in both genotypes. The latter would allow to directly relate the animal’s decision with AC activity patterns, which have been shown to correlate (Bathellier et al. 2012; Xin et al. 2019).

As α_2_δ3 KO mice can distinguish PTs normally from each other (Pirone et al. 2014), the lower local heterogeneity in frequency tuning observed in AAF and A2 appears to be of little consequence regarding perceptual or learning defects. Nevertheless, the observation suggests another developmental consequence of impaired temporal processing in the auditory brainstem. In general, AC tonotopy can be observed well on a global scale, but tuning is quite heterogeneous on a local scale (Bandyopadhyay et al. 2010; Rothschild et al. 2010; Romero et al. 2019; Tischbirek et al. 2019; Gaucher et al. 2020) which raises the question whether this is just a result of imprecision within the topographic network, or the expression of AC topography being governed by other sound aspects as well, which are not taken into account during BF mapping. Impaired temporal processing of sounds in α_2_δ3 KO mice might thus lead to an overemphasis of specifically the frequency aspect of sounds perceived during development, resulting in a slightly improved tonotopy within the AC. Our observation of a larger amount of well-tuned neurons in AAF and A2 is in line with this. However, as stated above, the generally less heterogeneous distribution of neurons’ BF in these two subfields might also be the cause of the lower local heterogeneity. At this point, it remains unknown why local tuning heterogeneity is unchanged, or even slightly elevated in A1, but it is in agreement with the idea that different AC subfields fulfill different functions in sound processing, and thus may react differentially to central impairments. In the future, it would be interesting to study synaptic connections within and to the AC in α_2_δ3 KO mice to decipher the direct cause of the observed aberrations. For example, neurons with similar orientation selectivity in the visual cortex have an overproportionally high chance of being synaptically connected (Ko et al. 2011), and thus studying synaptic connectivity within the AC of α_2_δ3 KO mice might provide more insight into the cause of altered local tuning heterogeneity. In fact, α_2_δ3 KO mice exhibit alterations in axonal and dendritic processes in somatosensory cortex (Landmann et al. 2018), providing evidence for disturbed cortical connectivity. Furthermore, presynaptic defects, as discussed above for synapses within auditory brainstem, might obviously also be present in the AC of α_2_δ3 KO mice and contribute to the observed sound processing defects. It would thus be highly interesting to in the future use conditional α_2_δ3 KO mice to assess which of the alterations are caused by developmental defects, and which are direct consequences of altered sound-evoked activity within the auditory system. α_2_δ subunit defects are associated with a variety of brain disorders (Ablinger et al. 2020), and specifically *Cacna2d3* mutations have been identified as risk mutations for ASD in humans (Iossifov et al. 2012; Girirajan et al. 2013; De Rubeis et al. 2014). This neuropsychiatric disorder is most often associated with social and cognitive challenges, yet is also accompanied by pronounced sensory processing deficits (reviewed in Robertson and Baron-Cohen 2017), including the temporal auditory domain (Kwakye et al. 2011). Animal models of ASD have provided insight into impaired auditory processing, for example concerning cortical plasticity (Kim et al. 2013; Yang et al. 2014), auditory-related learning (Reinhard et al. 2019; Lee et al. 2022), and pitch discrimination (Rendall et al. 2019). However, to our knowledge our study is the first the first to investigate large-scale activity patterns in the AC of an ASD-associated mouse model employing two-photon activity imaging. Our results of altered network activity patterns and tuning topography in α_2_δ3 KO mice demonstrate the ability of this technique to provide new insight into central auditory processing defects and add to the understanding of activity alterations in the auditory system in ASD.

## Materials and methods

A detailed description can be found online as supplementary material.

### Animals

α_2_δ3 KO mice were generated by Deltagen (Neely et al. 2010) and purchased through The Jackson Laboratories (B6N(Cg)-*Cacna2d3*^*tm1b(KOMP)Wtsi*^/J, Bar Harbor, ME, USA). Gene function was corrupted by inserting a LacZ cassette into exon 15 (of 39 exons) thereby obtaining a lacZ reporter function (no indication for a smaller gene product exists). They were bred and housed at the animal facility of the Center for Integrative Physiology and Molecular Medicine at the University of Saarland, Homburg, and transported to the University of Kaiserslautern for experiments. All experiments were conducted in accordance with the German Animal Protection Law (§4, Absatz 3 and TschG §7, Absatz 2). In vivo experiments were approved by the animal welfare council of Rhineland-Palatinate under file number G19-2-032. Experiments were performed up to P70 (P50 to P70 for electrophysiological experiments). Mice of both genders were used, 3 per group for in vivo experiments.

### Injection of viral vectors, habituation, and window implantation

To provide analgesia, mice were injected with carprofen intraperitoneally prior to the initial anesthesia with isoflurane, and locally with lidocaine. After opening of the scalp, a small hole was drilled above the AC. The viral vector AAV1-hSyn-GCaMP7f (pGP-AAV-syn-jGCaMP7f-WPRE was a gift from Douglas Kim & GENIE Project (Addgene viral prep # 104488-AAV1)) was injected and a titanium anchor was attached using dental cement (C&B Metabond; Parkell Inc., Farmingdale, NW, USA). The remaining skin was cemented to the head plate, sealing the operation site. The animal was then brought to its home cage for recovery. For analgesia and to prevent inflammation, Carprofen was administered for two subsequent days.

Before performing imaging experiments in awake mice, animals were first habituated to the experimental situation, head fixation and treadmill walking. 8 days after AAV injection, a window was implanted into the skull. A round piece of skull over the AC, as well as the dura mater, were removed. A stack of round cover glass was lowered onto the brain and then fixed with dental cement.

### In vivo Ca^2+^-imaging

The cranial window was covered with ultrasound gel (Anagel, Ana Wiz Ltd, Addlestone, UK) for recordings with a 10x water-immersion objective (IMPPLFLN, Olympus K.K., Shinjuku, Japan) or 16x water-immersion objective (CFI75 LWD, 0.8 NA, Nikon, Tokyo, Japan). As excitation source for two-photon imaging a Chameleon Vision II Titan:Saphir-Laser (Coherent Inc. Santa Clara, CA, USA) was used, tuned to 940 nm. Imaging was performed using a Galvo-Resonant 8 kHz scanning microscope (Ultima Investigator, Bruker, Billerica, MA, USA), at 29.76 frames/s. Emitted fluorescence was collected by the 16x objective and guided through a green emission band-pass filter onto a GaAsP photomultiplier tube (Bruker). For widefield imaging, illumination with blue light was achieved by a LED (470 nm, Thorlabs GmbH, Bergkirchen, Germany). Emitted fluorescence was passed through a green emission filter onto a scientific CMOS camera (Prime 95B, Teledyne Photometrics, Tucson, AZ, USA).

### Sound stimulation

The light box as well as the microscope and treadmill position were covered with sound attenuating foam (Basotect®, BASF, Ludwigshafen, Germany) protecting the recording site from external background noise as well as scanner noise, respectively. The speaker was calibrated daily. During two-photon imaging two groups of acoustic stimuli consisted of 17 different PTs (4-64 kHz, 4 equivalent steps per octave; 250 ms tone length followed by a 1 s pause), AM tones (same carrier frequencies and time course as PTs, 20 & 40 Hz modulation frequency, 70 dB SPL), and complex acoustic stimulations, which consisted of 10 animal vocalizations (birds, bats, and insects, 1 s pause between vocalizations, 70 dB SPL; all vocalizations downloaded from “http://www.avisoft.com/animal-sounds/“). We thank Matthias Göttsche (Stocksee, Germany) for permitting the use of recordings of the Blasius’s Horseshoe Bat. For widefield imaging 5 PTs were randomly presented during each of 16 repetitions and played at 50, 60 and 70 dB SPL.

### Analysis of in vivo data

For analysis of widefield imaging data, the procedure of image processing was adapted from Romero et al. (2019). The frequency eliciting the highest response amplitude within a pixel was set as the BF of that given pixel. A vector-based calculation of reversal points was provided to assist subfield parcellation. For two-photon data, recordings were processed with suite2p (https://suite2p.readthedocs.io/ (Pachitariu et al. 2017)). ROI fluorescence traces, subtracted by 0.7 times neuropil traces, were then deconvolved using the OASIS algorithm (Friedrich et al. 2017) and resulting spiking probabilities were used for most of later analyses. The data were then exported to MATLAB for further processing.

Phases of running which exceeded 1 cm/s were excluded from the activity traces for analysis. Neurons were defined as PT-responsive by comparing the mean spiking probabilities 400 ms pre stimulus and 400 ms post stimulus onset. A one-way ANOVA compared both distributions for each frequency-intensity distribution and if p < 0.01 in at least one PT-SPL combination, neurons were classified as PT-responsive, others were excluded. Mean post stimulus onset responses of each frequency were averaged across SPLs resulting in a single mean value per frequency. These points were then fitted with a unimodal and bimodal gaussian fit function to check for single- or double-peak tuning, respectively. In each FRA, the frequency which elicited the highest response was defined as the BF, regardless of SPL. To find neurons with similar activity patterns upon animal vocalization or PT/AM sound stimulation, a correlation analysis was performed related to Bathellier et al., (2012).

### Acute slice physiology

Animals were anesthetized with isoflurane and subsequently decapitated. 270 µm thick coronal slices were prepared using a vibratome (Leica VT 1200S, Leica, Wetzlar, Germany) containing ice-cold NMDG-based preparation solution. For recovery, slices were then transferred to 37°C and after 7 min incubation stored at RT in artificial cerebral spinal fluid (aCSF).

Electrophysiological recordings, accompanied by single cell Ca^2+^ imaging were performed on an electrophysiological rig, equipped with differential interference contrast optics, appropriate objectives (Nikon 4x CFI Achromat, 0.1 NA; Nikon 60x CFI Fluor W, 1.0 NA) and a scientific CMOS camera (Iris 9, Teledyne Photometrics), controlled by Micro-Manager (v2.0). A blue light LED (470 nm, Thorlabs) was used for illumination. Whole-cell recordings were obtained using a patch-clamp amplifier (EPC9, HEKA Electronics, Lambrecht, Germany) and a micromanipulator (SM-I, Luigs & Neumann) linked to the head stage.

## Acknowledgments

We thank the BioComp Research Initiative for funding. This work was supported by Deutsche Forschungsgemeinschaft (DFG) PP1608 (En 294/5-2 to JE, KU 1972/5-2 to SK) and DFG SFB 894 (A8 to JE). We thank Matthias Göttsche (Stocksee, Germany) for permitting the use of the recordings of the Blasius’s Horseshoe Bat. We thank Kerstin Fischer (Saarland University) and Kornelia Ociepka (University of Kaiserslautern) for excellent technical assistance.

