## Supplementary figures, tables, and Materials and Methods for "Altered population activity and local tuning heterogeneity in auditory cortex of *Cacna2d3*-deficient mice"

### Supplementary Material

#### Supplementary Figures

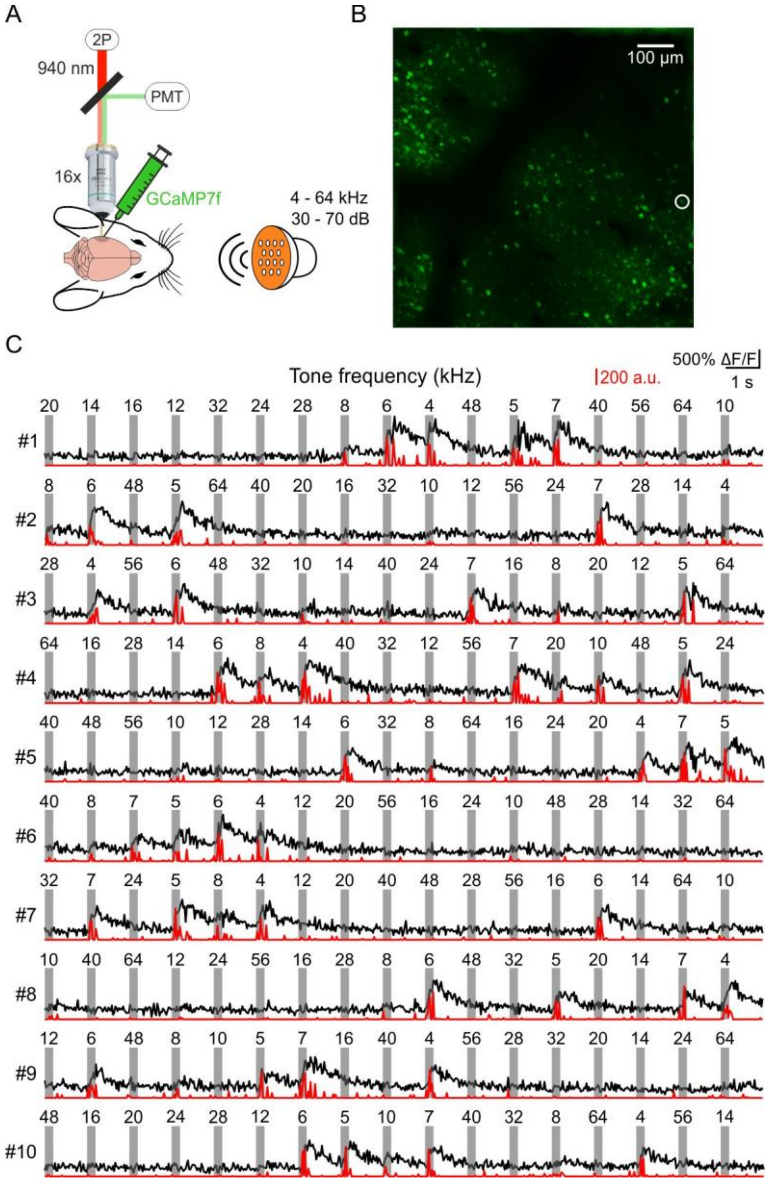

🡐 **Supplementary Figure 1**: In vivo awake 2-photon $Ca^{2+}$ imaging during PT stimulation in a $\alpha_2\delta 3$ KO mouse. (A) Scheme depicting the principle of imaging $Ca^{2+}$ activity of GCaMP7f-expressing neurons during PT stimulation via two-photon imaging. (B) Maximum intensity projection of a $Ca^{2+}$ activity time series from a $\alpha_2\delta 3$ KO FOV, recorded during PT stimulation at 60 dB SPL, 230 µm below the pial surface. (C) Uncut extracted fluorescent activity (ΔF/F0, black trace) from the neuron encircled in B (white circle) and corresponding deconvolution (red trace) during PT presentation at 60 dB SPL. Timings and length (250 ms) of presented PTs are depicted by grey bars with the corresponding frequency depicted above. Each row shows one repetition containing each of the 17 PTs once.

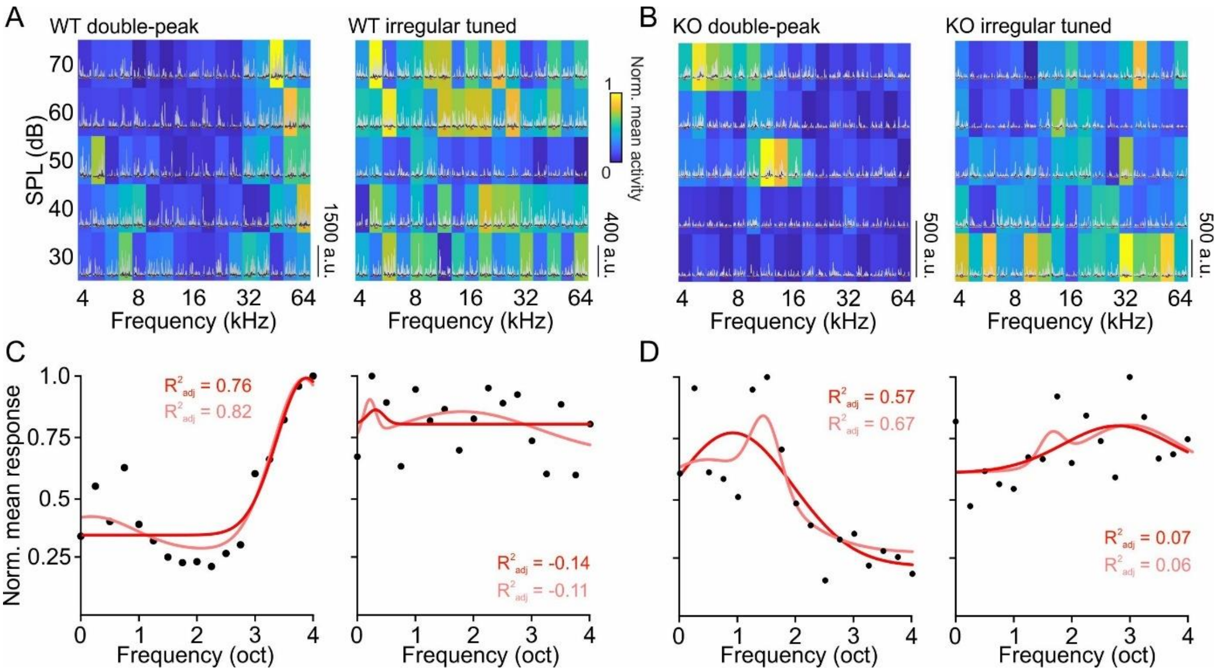


**Supplementary Figure 2**: FRAs of double-peak and irregular-tuned neurons. See legend of Figure 2A for details. (A) FRAs for WTs. (B) FRAs for KO. (C) Averaged activity across SPLs shown in A for each frequency (black dots). Unimodal (red) and bimodal (light red) gaussian fits. (D) As C, but for FRA in B.

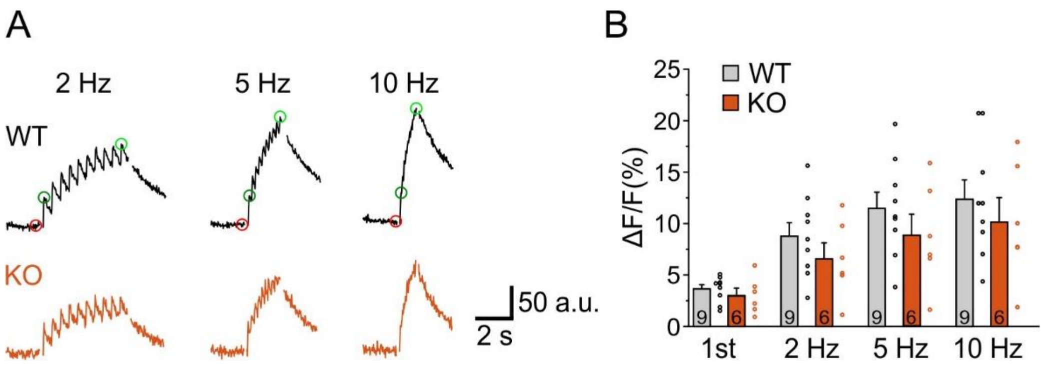


**Supplementary Figure 3**: $Ca^{2+}$ imaging control experiments. (A) Representative fluorescence traces of single OGB-filled neurons during AP-induction at different frequencies. Red circle depicts baseline value, green circles 1st AP and summation peak values. Gaps in the traces are due to dropped frames in recordings for technical reasons. (B) Statistics for fluorescence peaks as depicted in A. Numbers in bar depict number of neurons analyzed.

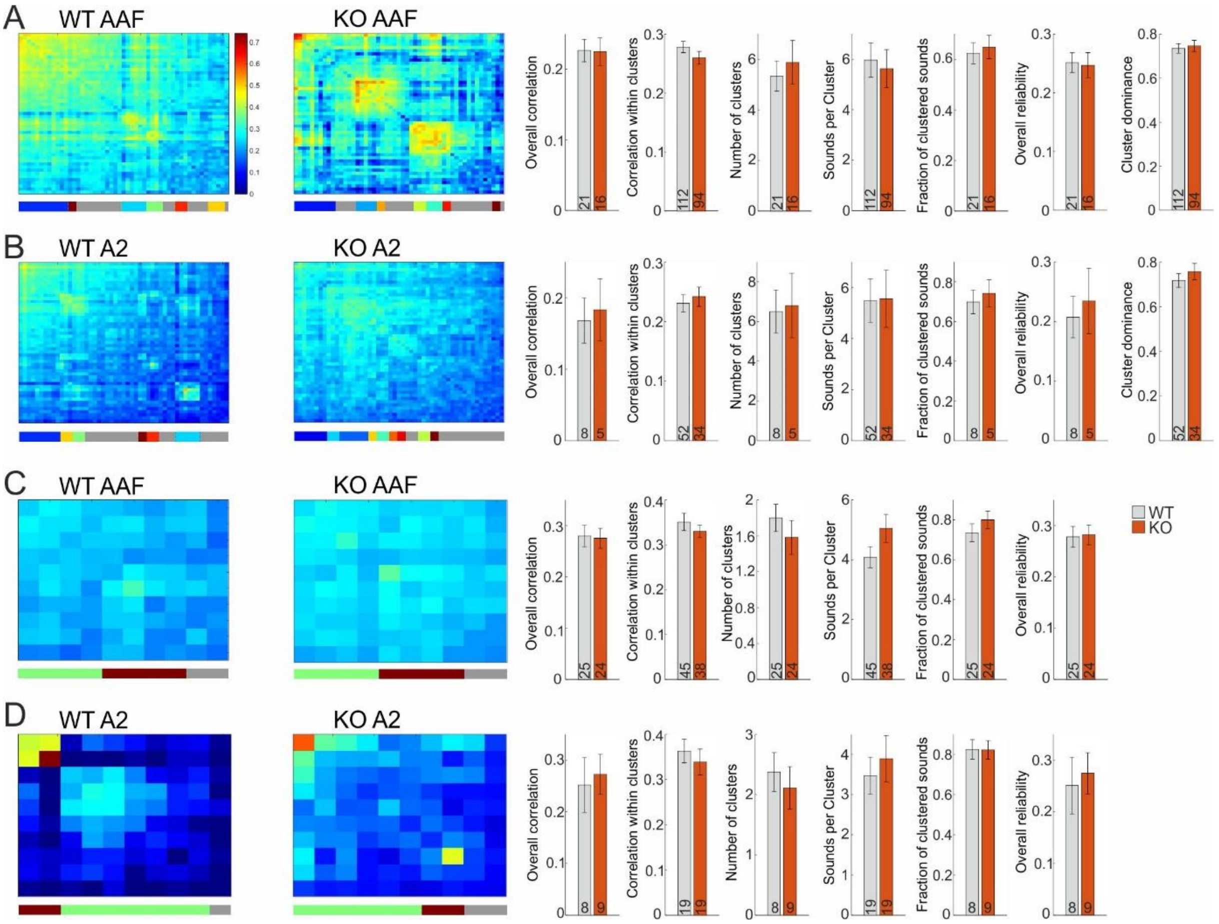


**Supplementary Figure 4**: Cluster analysis of sound-evoked activity in AAF and A2. (A) Left: Correlation matrix of 51 sound-evoked patterns (17 patterns for each of PTs, AM tones with 20Hz modulation, and AM tones with 40 Hz modulation) after hierarchical clustering, activity recorded in AAF. Vertical color bar at the bottom depicts clusters, grey stripes correspond to sounds that are not part of clusters. The diagonal depicts mean correlation across repetitions, thus showing reliability of the network. Right: Statistics for sound cluster analysis of PT/AM sounds. Numbers in bars depict n-number, either FOVs or sound clusters. (B) As A, but for A2. (C) Correlation matrix and statistics for 10 natural animal vocalizations, activity recorded in AAF. (D) As C, but for A2. See Supplementary Tables 1 and 2 for obtained order of sounds.

**Supplementary Tables**

**Supplementary Table 1:** Order of clustered PT/AM tones. Listed are the obtained orders of the exemplary correlation matrices shown in Figure 3A, and Supplementary Figure 4A and B.

| A1 WT | A1 KO | AAF WT | AAF KO | A2 WT | A2 KO |
|---|---|---|---|---|---|
| 7 kHz PT | 8 kHz AM 40 Hz | 8 kHz AM 20 Hz | 28 kHz PT | 10 kHz AM 20 Hz | 10 kHz PT |
| 7 kHz AM 40 Hz | 20 kHz AM 40 Hz | 7 kHz AM 40 Hz | 6 kHz AM 20 Hz | 12 kHz AM 20 Hz | 12 kHz PT |
| 5 kHz AM 40 Hz | 16 kHz AM 40 Hz | 6 kHz PT | 32 kHz PT | 28 kHz AM 20 Hz | 14 kHz PT |
| 5 kHz PT | 10 kHz AM 40 Hz | 40 kHz AM 40 Hz | 32 kHz AM 20 Hz | 64 kHz AM 40 Hz | 16 kHz PT |
| 20 kHz AM 20 Hz | 28 kHz AM 40 Hz | 40 kHz AM 20 Hz | 8 kHz AM 20 Hz | 24 kHz AM 40 Hz | 20 kHz PT |
| 10 kHz AM 20 Hz | 14 kHz AM 40 Hz | 7 kHz PT | 28 kHz AM 20 Hz | 5 kHz AM 20 Hz | 32 kHz PT |
| 8 kHz AM 20 Hz | 5 kHz AM 40 Hz | 48 kHz PT | 48 kHz AM 40 Hz | 14 kHz AM 20 Hz | 8 kHz PT |
| 4 kHz AM 20 Hz | 40 kHz AM 40 Hz | 16 kHz PT | 16 kHz AM 20 Hz | 16 kHz AM 40 Hz | 5 kHz PT |
| 16 kHz AM 20 Hz | 48 kHz AM 40 Hz | 4 kHz AM 40 Hz | 14 kHz AM 20 Hz | 48 kHz AM 40 Hz | 4 kHz PT |
| 14 kHz AM 20 Hz | 12 kHz AM 40 Hz | 20 kHz AM 40 Hz | 56 kHz AM 40 Hz | 5 kHz AM 40 Hz | 28 kHz PT |
| 5 kHz AM 20 Hz | 24 kHz AM 40 Hz | 24 kHz AM 40 Hz | 4 kHz PT | 6 kHz PT | 24 kHz PT |
| 7 kHz AM 20 Hz | 64 kHz AM 40 Hz | 28 kHz AM 40 Hz | 7 kHz AM 20 Hz | 7 kHz PT | 7 kHz PT |
| 28 kHz AM 40 Hz | 7 kHz AM 40 Hz | 24 kHz AM 20 Hz | 40 kHz AM 20 Hz | 8 kHz PT | 56 kHz PT |
| 32 kHz AM 20 Hz | 24 kHz AM 20 Hz | 8 kHz AM 40 Hz | 64 kHz AM 20 Hz | 12 kHz PT | 6 kHz PT |
| 6 kHz AM 20 Hz | 6 kHz PT | 32 kHz PT | 6 kHz PT | 16 kHz PT | 64 kHz PT |
| 24 kHz AM 20 Hz | 24 kHz PT | 8 kHz PT | 7 kHz PT | 64 kHz PT | 48 kHz PT |
| 40 kHz AM 20 Hz | 16 kHz PT | 28 kHz PT | 10 kHz AM 20 Hz | 48 kHz PT | 40 kHz PT |
| 28 kHz AM 20 Hz | 4 kHz AM 20 Hz | 40 kHz PT | 20 kHz PT | 40 kHz AM 20 Hz | 40 kHz AM 40 Hz |
| 10 kHz AM 40 Hz | 10 kHz PT | 56 kHz PT | 64 kHz PT | 12 kHz AM 40 Hz | 4 kHz AM 20 Hz |
| 4 kHz PT | 48 kHz PT | 32 kHz AM 20 Hz | 8 kHz PT | 56 kHz AM 20 Hz | 7 kHz AM 20 Hz |
| 10 kHz PT | 14 kHz PT | 12 kHz PT | 12 kHz PT | 8 kHz AM 20 Hz | 10 kHz AM 20 Hz |
| 12 kHz PT | 64 kHz AM 20 Hz | 6 kHz AM 40 Hz | 24 kHz PT | 16 kHz AM 20 Hz | 8 kHz AM 40 Hz |
| 14 kHz PT | 28 kHz AM 20 Hz | 64 kHz PT | 40 kHz PT | 20 kHz AM 40 Hz | 10 kHz AM 40 Hz |
| 56 kHz AM 20 Hz | 8 kHz PT | 16 kHz AM 40 Hz | 12 kHz AM 20 Hz | 8 kHz AM 40 Hz | 12 kHz AM 40 Hz |
| 24 kHz AM 40 Hz | 6 kHz AM 20 Hz | 10 kHz PT | 4 kHz AM 20 Hz | 64 kHz AM 20 Hz | 28 kHz AM 40 Hz |
| 28 kHz PT | 64 kHz PT | 4 kHz AM 20 Hz | 48 kHz AM 20 Hz | 28 kHz AM 40 Hz | 20 kHz AM 40 Hz |
| 32 kHz PT | 10 kHz AM 20 Hz | 6 kHz AM 20 Hz | 10 kHz PT | 32 kHz AM 40 Hz | 16 kHz AM 40 Hz |
| 64 kHz PT | 20 kHz AM 20 Hz | 12 kHz AM 20 Hz | 48 kHz PT | 48 kHz AM 20 Hz | 32 kHz AM 40 Hz |
| 24 kHz PT | 56 kHz PT | 48 kHz AM 20 Hz | 5 kHz PT | 24 kHz AM 20 Hz | 48 kHz AM 20 Hz |
| 12 kHz AM 40 Hz | 5 kHz AM 20 Hz | 10 kHz AM 20 Hz | 4 kHz AM 40 Hz | 4 kHz PT | 12 kHz AM 20 Hz |
| 48 kHz PT | 20 kHz PT | 32 kHz AM 40 Hz | 7 kHz AM 40 Hz | 5 kHz PT | 24 kHz AM 40 Hz |
| 56 kHz PT | 8 kHz AM 20 Hz | 16 kHz AM 20 Hz | 28 kHz AM 40 Hz | 32 kHz AM 20 Hz | 14 kHz AM 40 Hz |
| 8 kHz AM 40 Hz | 4 kHz AM 40 Hz | 20 kHz AM 20 Hz | 5 kHz AM 40 Hz | 6 kHz AM 40 Hz | 28 kHz AM 20 Hz |
| 40 kHz AM 40 Hz | 6 kHz AM 40 Hz | 56 kHz AM 20 Hz | 24 kHz AM 40 Hz | 7 kHz AM 40 Hz | 6 kHz AM 20 Hz |
| 48 kHz AM 40 Hz | 56 kHz AM 40 Hz | 28 kHz AM 20 Hz | 64 kHz AM 40 Hz | 4 kHz AM 20 Hz | 40 kHz AM 20 Hz |
| 20 kHz PT | 5 kHz PT | 14 kHz AM 20 Hz | 6 kHz AM 40 Hz | 56 kHz PT | 8 kHz AM 20 Hz |
| 20 kHz AM 40 Hz | 16 kHz AM 20 Hz | 64 kHz AM 40 Hz | 5 kHz AM 20 Hz | 10 kHz AM 40 Hz | 14 kHz AM 20 Hz |
| 16 kHz PT | 32 kHz AM 20 Hz | 5 kHz AM 40 Hz | 8 kHz AM 40 Hz | 56 kHz AM 40 Hz | 64 kHz AM 40 Hz |
| 48 kHz AM 20 Hz | 12 kHz AM 20 Hz | 5 kHz PT | 14 kHz AM 40 Hz | 10 kHz PT | 6 kHz AM 40 Hz |
| 8 kHz PT | 12 kHz PT | 14 kHz AM 40 Hz | 16 kHz PT | 14 kHz PT | 64 kHz AM 20 Hz |
| 40 kHz PT | 32 kHz PT | 56 kHz AM 40 Hz | 20 kHz AM 40 Hz | 20 kHz PT | 5 kHz AM 20 Hz |
| 6 kHz AM 40 Hz | 7 kHz PT | 48 kHz AM 40 Hz | 40 kHz AM 40 Hz | 24 kHz PT | 16 kHz AM 20 Hz |
| 14 kHz AM 40 Hz | 40 kHz AM 20 Hz | 20 kHz PT | 24 kHz AM 20 Hz | 28 kHz PT | 32 kHz AM 20 Hz |
| 16 kHz AM 40 Hz | 14 kHz AM 20 Hz | 12 kHz AM 40 Hz | 32 kHz AM 40 Hz | 32 kHz PT | 20 kHz AM 20 Hz |
| 56 kHz AM 40 Hz | 32 kHz AM 40 Hz | 10 kHz AM 40 Hz | 14 kHz PT | 6 kHz AM 20 Hz | 24 kHz AM 20 Hz |
| 32 kHz AM 40 Hz | 4 kHz PT | 5 kHz AM 20 Hz | 12 kHz AM 40 Hz | 20 kHz AM 20 Hz | 56 kHz AM 20 Hz |
| 4 kHz AM 40 Hz | 40 kHz PT | 4 kHz PT | 20 kHz AM 20 Hz | 40 kHz PT | 7 kHz AM 40 Hz |
| 6 kHz PT | 56 kHz AM 20 Hz | 14 kHz PT | 56 kHz AM 20 Hz | 14 kHz AM 40 Hz | 5 kHz AM 40 Hz |
| 64 kHz AM 40 Hz | 7 kHz AM 20 Hz | 7 kHz AM 20 Hz | 56 kHz PT | 40 kHz AM 40 Hz | 4 kHz AM 40 Hz |
| 12 kHz AM 20 Hz | 48 kHz AM 20 Hz | 64 kHz AM 20 Hz | 10 kHz AM 40 Hz | 7 kHz AM 20 Hz | 48 kHz AM 40 Hz |
| 64 kHz AM 20 Hz | 28 kHz PT | 24 kHz PT | 16 kHz AM 40 Hz | 4 kHz AM 40 Hz | 56 kHz AM 40 Hz |

**Supplementary Table 2:** Order of clustered natural animal vocalizations. Listed are the obtained orders of the exemplary correlation matrices shown in Figure 3C, and Supplementary Figure 4C and D.

| A1 WT | A1 KO | AAF WT | AAF KO | A2 WT | A2 KO |
|---|---|---|---|---|---|
| Great green bush cricket | Lesser field grasshopper | Blasius's horseshoe bat | Blasius's horseshoe bat | Blue tit | European water shrew |
| Meadow grasshopper | Vervain hummingbird | Lesser field grasshopper | European water shrew | Vervain hummingbird | Vervain hummingbird |
| Lesser field grasshopper | Great green bush cricket | Treecreeper | Vervain hummingbird | Blasius's horseshoe bat | Meadow grasshopper |
| Mottled grasshopper | Common noctule bat | Blue tit | Blue tit | European water shrew | Mottled grasshopper |
| Vervain hummingbird | Meadow grasshopper | European water shrew | Meadow grasshopper | Treecreeper | Treecreeper |
| Blasius's horseshoe bat | Mottled grasshopper | Great green bush cricket | Mottled grasshopper | Meadow grasshopper | Great green bush cricket |
| Treecreeper | Blasius's horseshoe bat | Mottled grasshopper | Lesser field grasshopper | Mottled grasshopper | Blue tit |
| Blue tit | Treecreeper | Meadow grasshopper | Common noctule bat | Lesser field grasshopper | Common noctule bat |
| European water shrew | Blue tit | Vervain hummingbird | Great green bush cricket | Common noctule bat | Blasius's horseshoe bat |
| Common noctule bat | European water shrew | Common noctule bat | Treecreeper | Great green bush cricket | Lesser field grasshopper |

#### Supplementary Materials and methods

##### Animals

Mice with a targeted deletion of the *Cacna2d3* gene coding for $\alpha_2\delta3$ with insertion of a bacterial β-galactosidase under its promoter (B6.129P2-Cacna2d3tm1Dgen) were generated by Deltagen (Neely et al. 2010) and purchased through The Jackson Laboratories (B6N(Cg)-*Cacna2d3*$^{tm1b(KOMP)Wtsi}$/J, Bar Harbor, ME, USA). The LacZ cassette was inserted into exon 15 (of 39 exons) thereby obtaining a lacZ reporter function (no indication for a smaller gene product exists). They were crossed on a C57Bl/6N background (Charles River, Sulzfeld, Germany) for at least 10 generations. They were bred and housed at the animal facility of the Center for Integrative Physiology and Molecular Medicine at the University of Saarland, Homburg, and transported to the University of Kaiserslautern for experiments (at least 7 days adaption time between transport and start of experiments). Food and water were provided *ad libitum,* and all animals were housed at a 12-hour light-dark cycle. All experiments were conducted in accordance with the German Animal Protection Law (TschG §4, Absatz 3 and §7, Absatz 2). In vivo experiments were approved by the animal welfare council of Rhineland-Palatinate under file number G19-2-032. Experiments were performed up to postnatal day (P) 70 (P50 to P70 for electrophysiological experiments). Mice of both genders were used, 3 per group for in vivo experiments. The experimenter was blind to the genotype during experiments and initial processing of data.

##### Injection of viral vectors, habituation, and window implantation

To provide analgesia, mice were injected with carprofen intraperitoneally prior to the initial anesthesia with isoflurane (3-5 % in $O_2$, Fluovac system, Harvard Apparatus, Holliston, MA, USA). The anesthetized mouse was placed on a heating matt (TC-1000 Temperature Controller, CWE, Inc., Ardmore, PA, USA) and fixed in a stereotactic frame (Model 900, David

Kopf Instruments). The head holder was connected to the Fluovac system which allowed continuous supply of isoflurane (1-3 % in $O_2$) during the whole operation. After the fur was shaved from the scalp, 50 µl of lidocaine was injected subcutaneously for local analgesia. After 5 min waiting time, the scalp was opened along the midline and removed over the right hemisphere. Afterwards, the head was tilted by 45° to the right and the skin over the left hemisphere was pushed aside to reveal the underlying skull and muscles. The *musculus temporalis* was then partly removed to access the skull area over the AC. The area of the AC was then approximated by topographic structures and two injection sites were chosen. A small hole was drilled with a dental drill. The tip of a syringe (NanoFil, World Precision Instruments LLC, Sarasota County, FL, USA) containing 3.8 x $10^{12}$ GC/ml of the viral vector AAV1-hSyn-GCaMP7f (pGP-AAV-syn-jGCaMP7f-WPRE was a gift from Douglas Kim & GENIE Project (Addgene viral prep # 104488-AAV1)) was inserted through the hole. 750 nl of vector solution was injected at a rate of 80 nl/min at 500 µm depth. After successful injection, the syringe was kept in place for 5 min and the procedure was subsequently repeated for the second injection site. A titanium anchor was attached using dental cement (C&B Metabond; Parkell Inc., Farmingdale, NW, USA). The remaining skin from the left side of the scalp was then again pulled above the injection sites and cemented to the head plate, sealing the operation site. The animal was then brought to its home cage for recovery for at least two days before the start of habituation procedure. For analgesia and to prevent inflammation, Carprofen was administered for two subsequent days.

Before performing imaging experiments in awake mice, animals were first habituated to the experimental situation. In each session, animals were brought to a treadmill (LN treadmill, Luigs & Neumann, Ratingen, Germany) under the imaging setup and were allowed to freely explore their surroundings for 15 min. Subsequent head fixation lasted initially 10 min, increased by 25-30 min each day until 2 hours were reached, resembling the maximal time for one imaging session. Animals were not habituated on the day of window implantation surgery and on the two following days. After 5 habituation sessions, none of the animals showed any signs of stress, e.g., cowering, clinging, and sudden fast running, and were therefore used for subsequent imaging experiments.

8 days after AAV injection, a window was implanted into the skull, following the general surgical procedures described above. The joint between cement and remaining skin over the left hemisphere was reopened. The boundary of a round piece of skull, ~3 mm in diameter, over the auditory cortex (AC) was thinned down by tracing the edge with a dental drill. Once the remaining skull encircled by the furrow was loose it was removed with a fine kinked probe. The dura mater was removed with a small needle and fine forceps. A stack of 2 x 3 mm round cover glass (thickness #0, Warner Instruments LLC, Hamden, CT, USA) with 1 x 4 mm round cover glass (thickness #1), glued together by UV-curing adhesive (NOA 60, Norland Optics, East Windsor, NJ, USA), was lowered onto the brain, sealing the opening in the skull. The stack was then fixed with dental cement.

**In vivo $Ca^{2+}$-imaging**

For awake in vivo $Ca^{2+}$ imaging, the cranial window was covered with ultrasound gel (Anagel, Ana Wiz Ltd, Addlestone, UK) for recordings with a 10x water-immersion objective (IMPPLFLN, Olympus K.K., Shinjuku, Japan) and 16x water-immersion objective (CFI75 LWD, 0.8 NA, Nikon, Tokyo, Japan). All recordings were obtained using the Ultima Investigator microscope (Bruker AXS SAS, France) with the objective tilted by 45°, maintaining the animal in an upright position. As excitation source for two-photon imaging a Chameleon Vision II Titan:Saphir-Laser (Coherent Inc. Santa Clara, CA, USA), controlled by Chameleon Vision (v2.83, Coherent), with 140 fs pulse duration and 80 MHz repetition rate, with built-in precompensation, was used, tuned to 940 nm. Laser intensity was adjusted with a Pockels cell (Model 302RM Driver, Conoptics Inc., Danbury, CT, USA) and was in the range between 13 - 65 mW. Imaging was performed using a Galvo-Resonant 8 kHz scanner, recording a 512 pixel x 512 pixel field of view (FOV), covering ~820 µm x 820 µm, at 29.76 frames/s. Emitted fluorescence was collected by the 16x objective and guided through a green emission band-pass filter onto a GaAsP photomultiplier tube (Bruker). All imaging components were controlled by Prairie View (v5.5.64.500, Bruker) and parameters were set by MATLAB 2020a via Prairie Link (v5.5.0.48, Bruker). During all recordings, the movement of the mouse on the treadmill was registered by two hall sensors. On the first day of imaging, the FOVs were chosen to cover as much area as possible of the cranial window, depending on the area of transfected tissue. Experiments were performed over several days (data analyzed here were recorded over up to 5 days), covering the extent of the AC as much as possible across all subfields and different depths in layer 2/3.

For widefield imaging, illumination with blue light was achieved by a LED (470 nm, Thorlabs GmbH, Bergkirchen, Germany). Emitted fluorescence was passed through a green emission filter onto a scientific CMOS camera (Prime 95B, Teledyne Photometrics, Tucson, AZ, USA), controlled by Micro-Manager (v2.0, "https://micro-manager.org"). The corresponding FOV size covered ~1.5 mm x 1.5 mm resulting in a 1024 pixel x 1024 pixel image. Data were collected, after focusing roughly 200 µm below the pial surface, at a rate of 10 Hz with an exposure time of 60 ms.

**Sound stimulation**

The light box as well as the microscope and treadmill position were covered with sound attenuating foam (Basotect®, BASF, Ludwigshafen, Germany) protecting the recording site from external background noise as well as scanner noise, respectively. With this, the sound pressure level (SPL) of ambient noise in the relevant hearing range for mice (1-90 kHz) and in the range of frequencies used for sound stimulation (4-100 kHz) approximated 35 dB SPL and 20 dB SPL at max, respectively. Ambient noise was recorded with a high sensitive microphone (378A06, 3 - 40000 Hz, 12.6 mV/Pa, inherent noise: 22 dB(A) re 20 µPa, PCB Piezotronics, Depew, NY, USA) and the corresponding signal was amplified by an analog amplifier (MA3, Tucker-Davis Technologies (TDT), Alachua, FL, USA), coupled with an analog to digital multifunction processor (RX6, TDT) controlled by SigCalRP (v4.2; TDT). Daily calibration of the speaker was done using the same equipment except for a less sensitive microphone, which could record higher frequencies up to 100 kHz (Model 378C01, 4 – 100000 Hz, 2.01 mV/Pa, inherent noise: 42 dB(A) re 20 µPa, PCB). The SPL during tone presentation (4 – 80 kHz) was recorded and used to adjust driving voltages of the speaker for each frequency according to the desired SPL. Those normalized values were then exported to Matlab and filter coefficients were calculated used for the online FIR filter of sound presentation during experiments. During two-photon imaging two groups of acoustic stimuli consisted of 17 different pure tones (PTs, 4-64 kHz, 4 equivalent steps per octave; 250 ms tone length followed by a 1 s pause), amplitude-modulated (AM) tones (same carrier frequencies and time course as PTs, 20 & 40 Hz modulation frequency, 70 dB SPL), and complex acoustic stimulations, which consisted of 10 animal vocalizations (birds, bats, and insects, 1 s pause between vocalizations, 70 dB SPL; all vocalizations downloaded from http://www.avisoft.com/animal-sounds/). We thank Matthias Göttsche (Stocksee, Germany) for allowing us to use the recordings of the Blasius's Horseshoe Bat. Each tone/vocalization was high-pass filtered at 4 kHz. They were randomly presented within the sound group (PTs, AM tones, or animal vocalizations) during each of 10 repetitions. For widefield imaging 5 PTs (4, 8, 16, 32, 64 kHz, 500 ms tone; 5 s pause between tones) were randomly presented during each of 16 repetitions and played at 50, 60, and 70 dB SPL. All stimuli were created with MATLAB 2020a (MathWorks, Natik, MA, USA) controlling a script written in RPvdsEX (v88, TDT) and loaded to the RX6 digital-to-analog converter. The output signal of the RX6 was passed by an electrostatic speaker driver (ED1, TDT) and finally transformed into acoustic signals by an electrostatic free field speaker (ES1, TDT), positioned ~10 cm from the ear of the mouse contralateral to the window.

**Analysis of in vivo data**

For analysis of widefield imaging data, the procedure of image processing was adapted from Romero et al. (2019). Raw images were downsampled to a 256 pixel x 256 pixel resolution. Small drifts in fluorescence signal were removed by computing a temporal baseline (F0) for each pixel from a polynomial fit (degree 3) of a 15 s sliding window (Chronux toolbox, Matlab). The change in fluorescence was calculated for each frame as percent change from the temporally smoothed signal ($\Delta F/F\_0 \cdot 100$). These amplitudes were used for further analysis. Baseline activity levels for each stimulus were defined for each pixel by creating a histogram of amplitudes of all frames during the 2 s prestimulus period. To check for tone evoked responses, the maximum amplitude was picked from the 750 ms poststimulus onset period and averaged with the preceding and following frame. In cases where the resulting value exceeded the prestimulus baseline activity distribution by at least 2 standard deviations (z-score > 2), the response was characterized as tone evoked. A frequency-specific response amplitude was only calculated when a tone-evoked response occurred in a minimum of 4 of 16 repetitions. The frequency-specific response was then calculated as the mean of all significant tone-evoked response amplitudes to the respective frequency. The frequency eliciting the highest mean response amplitude within a pixel was set as the best frequency (BF) of that given pixel. As one FOV acquired with the 10x objective covered only a part of the AC, multiple overlapping FOVs were necessary in order to create a gapless BF map. The frequency-specific response amplitudes of each pixel within a FOV were normalized to provide comparability. In cases where one pixel was represented more than once (due to overlapping FOVs), the BF with the higher normalized mean response amplitude was chosen. Next, a vector-based calculation of reversal points, similar to the analysis in Romero et al. (2019), was provided as follows to assist subfield parcellation. First, centers of existing low-frequency hubs were identified. From each of these a set of 1440 radial vectors from 0-360° (0.25° step size) were drawn. The mean BFs along each radial vector (±1°) were smoothed with a moving average (window size 10 frames). The smoothed values were then fitted with a gaussian filter (degree 3), so that reversal points (first maxima) could be marked in the BF map. The end of the AC was defined as the point, where 10 pixels in a row showed no sound-evoked response at all. The marked reversal and end points within the BF map served as a template for the "drawassist" function of Matlab. Thereby, the subfield borders could be drawn by hand, but the outline was corrected by the information of the underlying BF map. Assignment of primary auditory field, anterior auditory field, and secondary auditory field was performed, based on existing knowledge from earlier studies (Tsukano et al., 2015; Tsukano et al., 2016; Romero et al., 2019).

For two-photon data, recordings were processed with suite2p (https://suite2p.readthedocs.io/; (Pachitariu et al. 2017)), first correcting for rigid as well as non-rigid movement shifts. Next, region of interest (ROI) detection, signal extraction and local neuropil signal extraction were carried out. ROI fluorescence traces, subtracted by 0.7 times neuropil traces, were then deconvolved using the OASIS algorithm (Friedrich et al. 2017) and resulting spiking probabilities were used for most of later analyses. ROIs were grouped as "cell" or "non-cell" by a trained classifier depending on activity parameter and parameter of the ROI shape. This automatic classification of ROIs as cells or non-cells was manually reviewed. The data were then exported to MATLAB for further processing.

AC activity is influenced during movement by inhibiting neuronal activity (Nelson et al. 2013). Therefore, phases of running which exceeded 1 cm/s were excluded from the activity traces for analysis. Furthermore, unresponsive neurons were removed from analysis if their peak signal-to-noise ratio (PSNR, Equation 1) was below 36 dB for the whole activity trace.

$$PSNR = 20 * log_{10}(\frac{max\ (F_{raw} - F_n)}{\sigma_n})$$ Equation 1 – PSNR

With $F_{raw}$, $F_n$, and $\sigma_n$ as the ROI trace, neuropil trace and the standard deviation of the neuropil trace, respectively.

Neurons were defined as PT-responsive by comparing the mean spiking probabilities 400 ms pre stimulus and 400 ms post stimulus onset. A one-way ANOVA compared both distributions for each frequency-intensity distribution and if $p < 0.01$ in at least 1 PT-SPL combination, neurons were classified as PT-responsive, others were excluded. Mean post stimulus onset responses of each frequency were averaged across SPLs resulting in a single mean value per frequency. These points were then fitted with an unimodal and bimodal gaussian (Equation 2 and Equation 3, respectively) fit function to check for single- or double-peak tuning, respectively.

$$Gauss1 = A_1 * e^{-(\frac{x-B_1}{C_1})^2} + D$$ **Equation 2 – gauss1**

$$gauss2 = A_1 * e^{-(\frac{x-B_1}{C_1})^2} + A_2 * e^{-(\frac{x-B_2}{C_2})^2} + D$$ **Equation 3 – gauss2**

To adjust for the different number of parameters of the two fits, the adjusted coefficient of determination ($R^2_{adj}$) was used to decide which fit was more suitable. If both fits resulted in

$R2adj < 0.4$ the neuron was classified as “irregular-tuned”, otherwise the fit which resulted in a higher $R^2_{adj}$ was used to assign a “single-“ or “double-peak tuning”. Furthermore, the tuning BW of each peak was determined as the full width at half maximum (Equation 4).

$$BW = 2 * \sqrt{2 * log_{10}(2) * \frac{C_{1/2}}{\sqrt{2}}}$$ **Equation 4 – BW**

In each Frequency response area (FRA), the frequency which elicited the highest response was defined as the BF, regardless of SPL. Next, local FOV coordinates from all neurons were converted into a global coordinate system. Global coordinates from each neuron were used to calculate the local BF distribution. For each neuron, its BF and the BFs of all neurons within 100 µm were extracted. Then, the interquartile range (IQR) of the distribution was calculated as a measure of local heterogeneity. If less than 5 neurons were within 100 µm radius, no IQR was calculated. Center frequency (CeF) was defined as the peak of the single unimodal gaussian fit.

To find neurons with similar activity patterns upon animal vocalization or PT/AM sound stimulation, a correlation analysis was performed related to Bathellier et al. (2012). Hence, the activity of each neuron in a time window of 0.55 s (0.4 s in case of PT/AM) from start of acoustic stimulation was averaged and the resulting values were combined to a “cell vector”. Vectors were then correlated across sounds (using Pearson correlation), determining the similarity, and these correlation values were averaged across repetitions, describing the reliability of responses. Hierarchical ordering of the resulting correlation matrix was carried out using unweighted average distance. The correlation matrix was exported in R (R Language, v4.1.2, https://www.R-project.org/) where clusters were defined by a hierarchical cluster tree using dynamic tree cut (Langfelder et al. 2007, method “hybrid”, “deepsplit” set to 2.5). To exclude clusters resulting from randomly formed correlations, a shuffling algorithm was implemented, randomly permutating the cell vector before calculations, repeated 5 times per FOV. Sound clusters in the real dataset were excluded if their average correlation was not at least two standard deviations above the average of correlations of shuffled clusters for the FOV.

**Acute slice physiology**

Animals were anesthetized with isoflurane and subsequently decapitated. The head was immediately submerged in ice-cold NMDG-preparation solution (in mM: 93 NMDG, 30 $NaHCO_3$, 20 HEPES, 25 D(+)-glucose, 3 myo-inositol, 2.5 KCl, 3 Na-pyruvate, 2 $CaCl_2$, 10 $MgCl_2$, 1.2 $NaH_2PO_4$, 5 ascorbic acid, pH 7.4, bubbled with carbogen (5 % CO2/ 95 % O2)). The brain was removed and the forebrain region containing the AC was cut out. 270 µm thick coronal slices were prepared using a vibratome (Leica VT 1200S, Leica, Wetzlar, Germany) containing ice-cold NMDG-preparation solution. For recovery, slices were then transferred to a beaker containing 37°C NMDG-preparation solution and after 7 min incubation stored at RT in artificial cerebral spinal fluid (aCSF, in mM: 125 NaCl, 25 $NaHCO_3$, 10 D(+)-glucose, 3 myo-inositol, 2.5 KCl, 2 Na-pyruvate, 2 $CaCl_2$, 1 $MgCl_2$, 1.25 $NaH_2PO_4$, 0.44 ascorbic acid, pH 7.4, bubbled with carbogen) until they were used for electrophysiological experiments.

Electrophysiological recordings, accompanied by single cell $Ca^{2+}$ imaging, were performed on an electrophysiological rig. Acute brain slices containing the AC were transferred into a recording chamber and fixed with a U-shaped platinum-iridium grid stringed with nylon strands. The chamber was then mounted on an upright microscope (Eclipse E600FN, Nikon), equipped with differential interference contrast optics, appropriate objectives (Nikon 4x CFI Achromat, 0.1 NA; Nikon 60x CFI Fluor W, 1.0 NA) and a scientific CMOS camera (Iris 9, Teledyne Photometrics), controlled by Micro-Manager (v2.0). During imaging experiments, a blue light LED (470 nm, Thorlabs) was used to illuminate the whole slice (1.1 $mW/cm^2$ at maximum) and emitted fluorescence was guided through a bandpass filter (500 nm - 550 nm) and captured by the camera. Once transferred to the microscope, the slices were continuously perfused with aCSF (RT, pH 7.4, bubbled with carbogen) using a peristaltic pump (ISM796B, Ismatec, Opfikon, Schweiz). Pipettes pulled from borosilicate glass capillaries with filament (GB150F-8P, Science Products, Hofheim am Taunus, Germany) using a horizontal puller (Flaming Brown Micropipette Puller P-87, Sutter instruments, Novato, CA, USA) had resistances ranging from 2.5 MΩ to 4.3 MΩ when filled with internal solution (in mM: 140 K-gluconate, 10 HEPES, 1 MgCl2, 2 ATP-$Na_2$, 0.3 GTP-$Na_2$, 0.05 Oregon green BAPTA-1 (OGB-1)). Whole-cell recordings were obtained using a patch-clamp amplifier EPC9, HEKA Electronics, Lambrecht, Germany) and a micromanipulator (SM-I, Luigs & Neumann) linked to the head stage. Capacitive transients were neutralized and series resistance was compensated by 50 – 70 %. Liquid junction potential (15.4 mV) was corrected online. During the 15 min filling time of the cell with internal solution, parameters for action potential (AP) generation, i.e., intensity and duration of the injected current were determined. Injected currents ranged from 0.3 nA - 1 nA with a duration of 2 - 8 ms. Once a suitable concentration of OGB-1 was achieved, rectangular current injections with were performed at different frequencies, while simultaneously recording fluorescent responses. The protocols were repeated up to three times. The obtained electrophysiological data was digitized with a sampling rate of 50 kHz and low-pass filtered at 8.4 kHz. Imaging data was sampled at 40 Hz with an exposure time of 25 ms. Image sequences recorded from OGB-1-filled cells were analyzed using a custom written MATLAB GUI allowing to display electrophysiological and fluorescence traces in parallel. Baseline for ΔF/F values was calculated by taking the mean of 75 ms prior to the peak.

To normalize changes in fluorescence across animals and slices ΔF/F values were calculated by averaging fluorescence 330 ms prior to each peak yielding a local baseline, which was used to normalize the corresponding peak. Hence, ΔF/F values for each stimulation intensity could be obtained.

##### Data visualization and statistics

Bar graphs and values in text present mean ± standard error of mean with number in bar depicting n-number. Normal distribution was tested with Kolmogorov-Smirnov test. Normally distributed data sets were compared using unpaired, two-tailed t-tests. Distribution-free data sets were compared using Mann-Whitney U-tests. Significance levels are as follows: $p < 0.05$ *, $p < 0.01$ **, $p < 0.001$ ***
